# Genomic epidemiology of global *Klebsiella pneumoniae* carbapenemase (KPC)-producing *Escherichia coli*

**DOI:** 10.1101/111914

**Authors:** N Stoesser, AE Sheppard, G Peirano, LW Anson, L Pankhurst, R Sebra, HTT Phan, A Kasarskis, AJ Mathers, TEA Peto, P Brandford, MR Motyl, AS Walker, DW Crook, JD Pitout

**Affiliations:** Modernising Medical Microbiology Consortium, Nuffield Department of Medicine, John Radcliffe Hospital, University of Oxford, Oxford, United Kingdom; Division of Microbiology, Calgary Laboratory Services, Calgary, Alberta, Canada; Department of Pathology and Laboratory Medicine, University of Calgary, Alberta, Canada; Icahn Institute and Department of Genetics and Genomic Sciences, Icahn School of Medicine, Mount Sinai, New York, USA; National Institute for Health Research (NIHR) Health Protection Research Unit (NIHR HPRU) in Healthcare Associated Infections and Antimicrobial Resistance, University of Oxford; Division of Infectious Diseases and International Health, Department of Medicine, University of Virginia Health System, Charlottesville, Virginia, USA; Office of Hospital Epidemiology, University of Virginia Health System, Charlottesville, Virginia, USA; AstraZeneca Pharmaceuticals LP, Waltham, Massachusetts, USA; Clinical Microbiology, Merck and Co Inc., Rahway, New Jersey, USA; Department of Microbiology, Immunology and Infectious diseases, University of Calgary, Alberta, Canada; Snyder Institute for Chronic diseases, University of Calgary, Alberta, Canada; Department of Medical Microbiology, University of Pretoria, South Africa

**Keywords:** *E. coli*, carbapenemase, *bla*_KPC_, whole genome sequencing

## Abstract

The dissemination of carbapenem resistance in *Escherichia coli* has major implications for the management of common human infections. *bla_KPC_,* encoding a transmissible carbapenemase (KPC), has historically largely been associated with *Klebsiella pneumoniae,* a predominant plasmid (pKpQIL), and a specific transposable element (Tn*4401,* ~10kb). Here we characterize the genetic features of the emergence of *bla*_KPC_ in global *E. coli,* 2008-2013, using both long-and short-read whole genome sequencing.

Amongst 43/45 successfully sequenced *bla*_KPC_-*E. coli* strains, we identified high strain (n=21 sequence types, 18% of annotated genes in the core genome); plasmid (≥9 replicon types); and *bla*_KPC_-associated, mobile genetic element (MGE) diversity (50% not within complete Tn*4401* elements). We also found evidence of interspecies, regional and international plasmid spread. In several cases *bla*_KPC_ was found on high copy number, small Col-like plasmids, previously associated with horizontal transmission of resistance genes in the absence of antimicrobial selection pressures.

*E. coli* is a common human pathogen, but also a commensal in a multiple environmental and animal reservoirs, and easily transmissible. The association of *bla*_KPC_ with a range of MGEs previously linked to the successful spread of widely endemic resistance mechanisms (e.g. *bla_T_*_EM_, *bla*_CTX-M_) suggests that it is likely to become similarly prevalent.

## INTRODUCTION

Carbapenemases have emerged over the last 15 years as one of the most significant antimicrobial resistance threats in Enterobacteriaceae, many species of which are major human pathogens^1^. They are enzymes with broad-spectrum hydrolytic activity targeting most beta-lactams, and commonly associated with other resistance mechanisms producing cross-resistance to other antimicrobial classes^2^. The *Klebsiella pneumoniae* carbapenemase (KPC) enzyme, encoded by alleles of the *bla*_KPC_ gene, represents one of the five major carbapenemase families, others being the VIM, IMP and NDM metallo-beta-lactamases, and the OXA-48-like oxacillinases^3^. The first KPC-producer, a *K. pneumoniae* strain harbouring *bla*_KPC-2_, was identified in 1996 in the eastern USA; since then, KPC-2 and KPC-3 (H272Y [C814T] with respect to KPC-2) have become widespread, and particularly entrenched in endemic hotspots in the USA, Greece, Israel, China and parts of Latin America^4,5^. The epidemic *K. pneumoniae* lineage, ST258, is thought to have contributed significantly to the global dissemination of *bla*_KPC-2_/*bla*_KPC-3_^6^, although these genes have now been observed in several species in the family Enterobacteriaceae^7,8^.

Acquired carbapenem resistance in *Escherichia coli* was considered rare as recently as 2010, although the first cases of *bla*_KPC_ in *E. coli* were observed as early as 2004-2005 in Cleveland (n=1, KPC-2^9^), New York City (n=2, KPC-2), New Jersey, USA (n=1, KPC-3)^10^, and Tel Aviv, Israel (n=4, KPC-2)^11,12^. No apparent epidemiological links were observed between any of these cases. Genotyping was limited at this time, but supported diversity being present in both host *E. coli* and *bla*_KPC_ plasmid backgrounds. Since then, direct, plasmid-mediated transfer of *bla*_KPC_ into *E. coli* within human hosts has been observed^13^, and clusters of KPC-producing *E. coli* have been identified in several geographic locations, from China to Puerto Rico^14,15^, and in the context of clinical infections^14,15^, asymptomatic colonization^16^ and in environmental isolates^17^.

More recently there has been significant concern around the identification of *bla*_KPC_ in *E. coli* sequence type (ST) 131, a globally disseminated and clinically successful strain^18-20^. Notably, the H30R/C1 clade (fluoroquinolone-resistant) and H30Rx/C2 clade (fluoroquinolone and extended-spectrum cephalosporin-resistant) sub-lineages of this strain have previously expanded globally in association with particular drug resistance mechanisms, including the extended-spectrum beta-lactamase (ESBL) gene, *bla*_CTX-M-15_ (clade C2)^21,22^. Given the high rates of community and healthcare-associated infections attributable to ST131^23^, and its capacity to be harbored asymptomatically in the gastrointestinal tract^24^, a stable association of ST131 with *bla*_KPC_ could have dramatic consequences for the management of *E. coli* infections^12^.

Despite these concerns, there are very limited detailed molecular epidemiological data investigating the genetic structures associated with *bla*_KPC_ in *E. coli* and the extent to which these may have been shared with other Enterobacteriaceae. Here we used a combination of short-read (Illumina) and long-read (PacBio) sequencing to reconstruct the chromosome and plasmid sequences of 44 *bla*_KPC_-positive *E. coli* isolates obtained consecutively from global surveillance schemes (67 participating countries, 2008-2013), fully resolving the *bla*_KPC_-containing plasmids in 24 cases, and comparing these data with other publicly available *bla*_KPC_ plasmid sequences.

## RESULTS

### Global *bla*_KPC_-*E. coli* strains are diverse, even within the most prevalent ST, ST131, with evidence for local transmission

45 isolates were obtained from 21 cities in 11 countries across four continents (2010-2013; previous laboratory typing results summarized in Supplementary Table 1). One isolate was *bla*_KPC_ negative on sequencing (ecol_252), and for one isolate the whole genome sequencing (WGS) data were inconsistent with the lab typing results on two occasions (ecol_451); these isolates were therefore excluded from analyses. Twenty-one different *E. coli* STs were represented amongst the remaining 43 isolates (Table 1; predicted in silico from whole genome sequencing), including: ST131 [n=16], ST410 [n=4], ST38 [n=3], ST10, ST69 [n=2 each] (remaining isolates singleton STs).

**Table 1.**
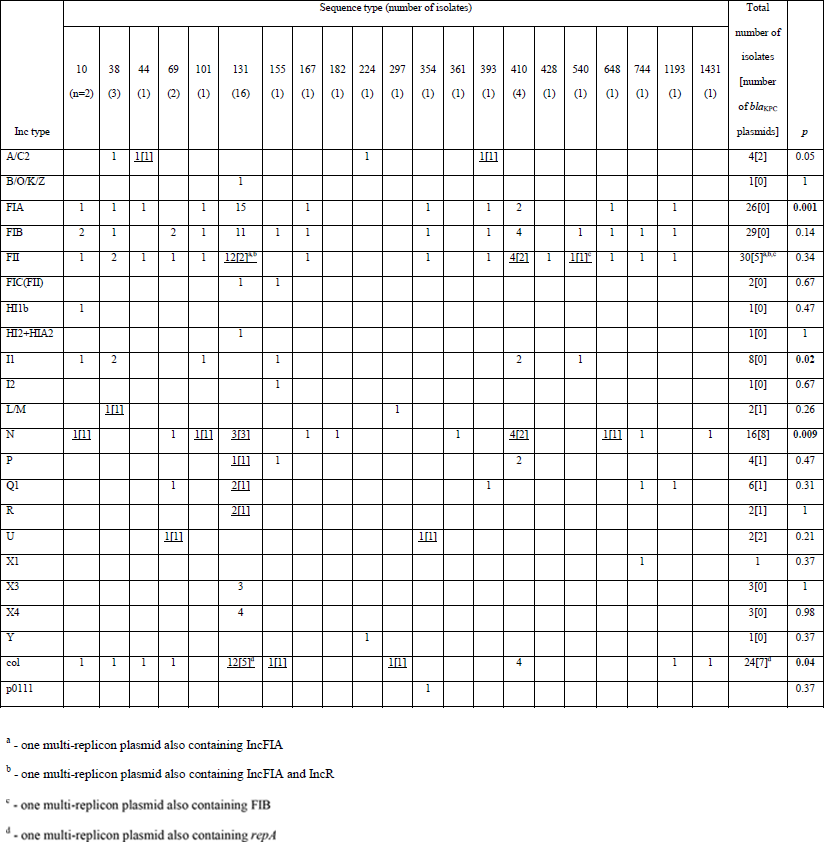
Plasmid replicon families present by ST, using the PlasmidFinder database^55^. Underlined text highlight categories associated with *bla*_KPC_; numbers in square brackets represent the known subset of *bla*_KPC_ plasmids in each cell. Exact test compares presence/absence of each Inc type by ST. The replicon type associated with *bla*_KPC_ could not be evaluated in 16 isolates, due to limitations of the assemblies.

Of 16,053 annotated open reading frames (ORFs) identified across all KPC *E. coli* isolates, only 2,950 (18.4%) were shared in all isolates (“core”), and a further 222 (1.4%) in 95-<100% of isolates (“soft core”^25^). At the nucleotide level there were 213,352 single nucleotide variants (SNVs) in the core genome, consistent with the previously observed diversity in the species^26^. Resistance gene profiles also varied markedly between strains, with some harbouring several beta-lactam, aminoglycoside, tetracycline and fluoroquinolone resistance mechanisms (e.g. ecol_224) and others containing *bla*_KPC_ only (e.g. ecol_584; Figure 1). For the 16 KPC-ST131 strains, 4,071/7,910 (50%) ORFs were core, with 6,778 SNVs across the core genome of these isolates, again consistent with previous global studies of ST131 diversity^21,22^ (Supplementary Figure S1). Accessory genomes were highly concordant for some (e.g. ecol_356/ecol_276/ecol_875), but not all (e.g. ecol_AZ159/ecol_244) isolates that were closely related in their core genomes, supporting highly variable evolutionary dynamics between core and accessory genomes (Figure 1). The geographic distribution of isolates closely related in both the core and accessory genomes supports local (e.g. ecol_AZ166, ecol_AZ167 [ST131, Beijing, China]) transmission of particular KPC-*E*. *coli* strains. The homology of genetic flanking motifs around the *bla*_KPC_ genes in these closely related isolate pairs would also be consistent with this hypothesis, and less consistent with multiple acquisition events of *bla*_KPC_ within the same genetic background, especially given the diversity in *bla*_KPC_ flanking sequences observed across the rest of the dataset (see below).

**Figure 1.**
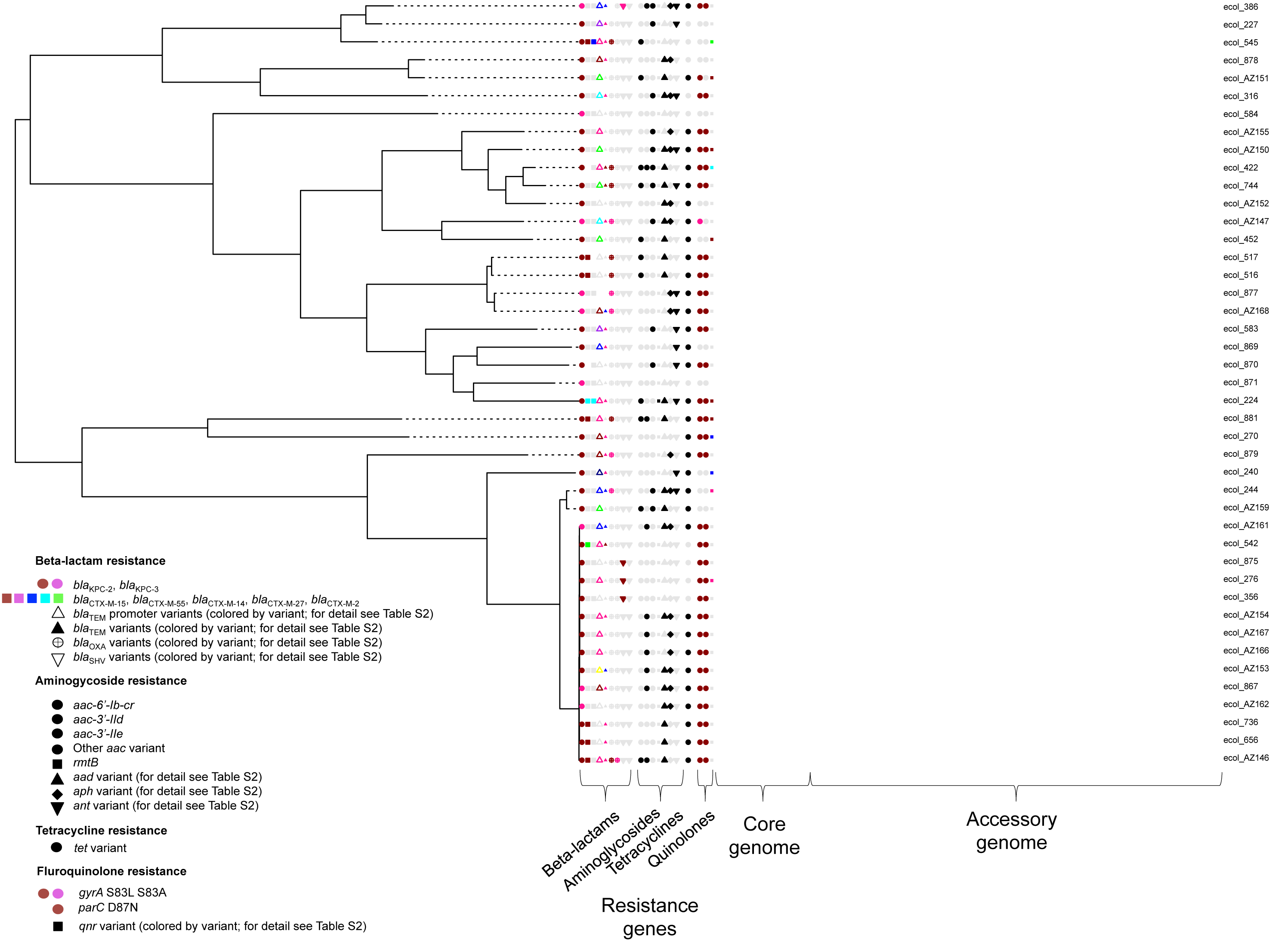
Phylogeny of KPC-*Escherichia coli* identified from global carbapenem resistance surveillance schemes, 2008-2013. Panels to the right of the phylogeny represent common resistance gene mechanisms (full details of resistance gene typing in Supplementary table 2), core and accessory genome components. For the accessory genome panel, blue represents annotated regions that are present, and grey those that are absent.

### *bla*_KPC_ genes appear restricted to plasmid contexts in *E. coli*, but may exist in multiple copies on single plasmid structures or in high copy number plasmids

Thirty-four isolates (80%) contained *bla*_KPC-2_, and nine isolates (20%) *bla*_KPC-3_. Chromosomal integration of *bla*_KPC_ has been previously described in other Enterobacteriaceae, *Pseudomonas* and *Acinetobacter* spp. but remains rare^8,27,28^; there was no evidence of chromosomal integration of *bla*_KPC_ in either the 18 chromosomal structures reconstructed from long-read sequencing or in *bla*_KPC_-containing contigs from all evaluable 42 isolates. *bla*_KPC_ alleles were not segregated by ST.

Estimates of *bla*_KPC_ copy number per bacterial chromosome varied between <1 (ecol_879, ecol_881) and 55 (ecol_AZ152). In nine cases this estimate was ≥10 copies of *bla*_KPC_ per bacterial chromosome (ecol_276, ecol_356, ecol_867, ecol_869, ecol_870, ecol_875, ecol_AZ150, ecol_AZ152, ecol_AZ159, Supplementary Table 2). Six of these isolates contained *bla*_KPC_ in a col-like plasmid context, in two cases the plasmid rep type was unknown, and in one case it was an IncN replicon. Plasmid copy number is associated with higher levels of antibiotic resistance if the relevant gene is located on a high-copy unit. Interestingly, high copy number plasmids are postulated to have higher chances of fixing in descendant cells, as they distribute more adequately by chance and without the requirement for partitioning systems^29^, and of being transferred in any conjugation event, either directly or indirectly^30-32^.

### *bla*_KPC_ and non- *bla*_KPC_ plasmid populations across global *bla*_KPC_*-E.coli* strains are extremely diverse

Plasmid Inc typing across all isolates revealed the presence of a median of four plasmid replicon types per isolate (range: 1-6; IQR: 3-5), representing wide diversity (Table 1). However, IncN, col, IncFIA and IncI1 replicons were disproportionately over-represented in certain STs (p<0.05; Table 1). Within the 18 isolates that underwent PacBio sequencing, we identified 53 closed, *non-bla*_KPC_ plasmids, ranging from 1,459 bp to 289,903 bp (Supplementary Table 1; at least four additional, partially complete plasmid structures were present). Of these non-*bla*_KPC_ plasmids, 10 (size: 2,571-150,994 bp) had <70% similarity (defined by percent sequence identity multiplied by proportion of query length demonstrating homology) to other sequences available in GenBank, highlighting that a significant proportion of the “plasmidome” in KPC-*E*. *coli* remains incompletely characterized. For the other 43 plasmids, the top match in GenBank was a plasmid from *E. coli* in 35 cases, *K. pneumoniae* in 5 cases, and *Citrobacter freundii, Shigella sonnei, Salmonella enterica* in 1 case each (Supplementary Table 3).

Twenty-three *bla*_KPC_ plasmid structures were fully resolved (five from Illumina data), ranging from 14,029 bp to 287,067 bp (median = 55,434 bp; IQR: 44,320-85,865 bp). These *bla*_KPC_-containing plasmids, and five additional cases where *bla*_KPC_ was identified on a replicon-containing contig, were highly diverse based on Inc typing (Supplementary Table 1). IncN was the most common type (n=7; 30%), followed by small, col-like plasmids (n=5 [col-like plasmids with single replicons only]; 22%). Other less common types were: A/C2, FII(k), U (all n=2); and L/M, P, Q1 and R (all n=1). Four (14%) *bla*_KPC_ plasmids were multi-replicon constructs, namely: *col/repA,* FIB/FII, FIA/FII, and FIA/FII/R.

### Common IncN plasmid backbones have dispersed globally within *E. coli*

From GenBank, we selected all unique, fully sequenced IncN-*bla*_KPC_ plasmid sequences (Supplementary Table 4) for comparison, dating from as early as 2005, around the time of the earliest reports of KPC-producing *E. coli.* The plasmid backbones and flanking sequences surrounding *bla*_KPC_ in these 16 plasmid references and a subset of 12 study sequences (see “Methods”) were consistent with multiple acquisitions of two known IncN-Tn*4401*-*bla*_KPC_ complexes in divergent *E. coli* STs: firstly, within a Plasmid-9 (FJ223607, 2005, USA)-like background, and secondly, within a Tn2/*3*-like element in a Plasmid-12 (FJ223605, 2005, USA)-like background.

In the first instance, genetic similarities were identified between Plasmid-9, pKPC-FCF/3SP, pKPC-FCF13/05, pCF8698, pKP1433 (representing a hybrid IncN), and *bla*_KPC_ plasmids from isolates ecol_516, ecol_517, ecol_656, and ecol_736 (this study). Plasmid-9 contains duplicate Tn*4401*b elements in reverse orientation with four different 5bp flanking sequences in an atypical arrangement within a group II intron^33^. The backbone structures of the other plasmids in this group are consistent with a separate acquisition event of a Tn*4401*b element between the *pld* and *traG* regions within an ancestral version of the Plasmid-9 structure, with the generation of a flanking TTCAG target site duplication (TSD) (labelled as Plasmid 9-like plasmid (hypothetical), Figure 2). International spread followed by local evolution both within and across species would account for the differences between plasmids, including: (i) nucleotide level variation (observed in all plasmids); (ii) small insertion/deletion events (observed in all plasmids); (iii) larger insertion/deletion events mediated by transposable elements (e.g. pCF8698_KPC_2); and (iv) likely homologous recombination, resulting in clustered variation within a similar plasmid backbone (e.g. ecol_656/ecol_736), as well as more distinct rearrangements, including the formation of “hybrid” plasmids (e.g. pKP1433)(Figure 2).

**Figure 2.**
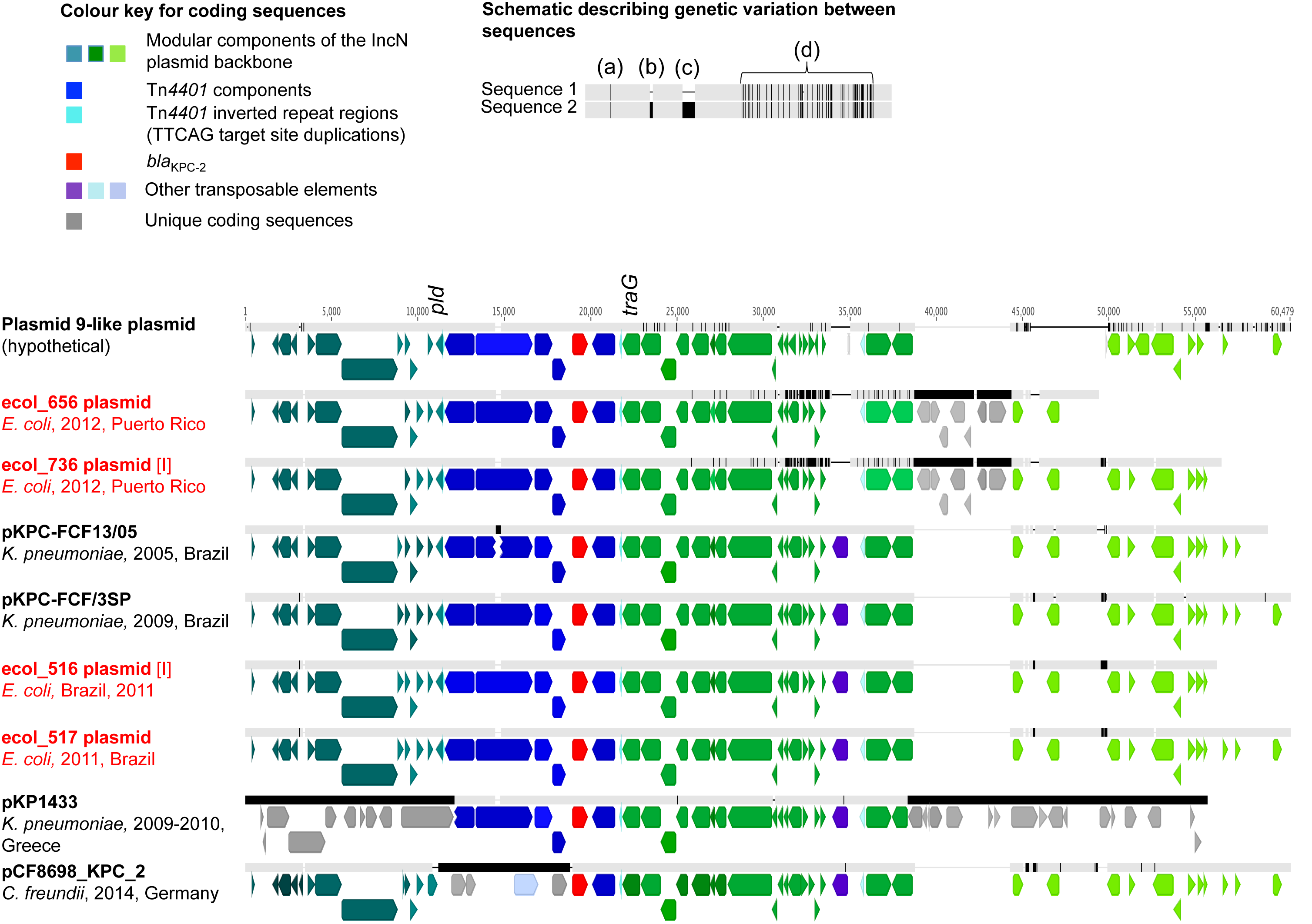
Comparison of FJ223607-like (Plasmid 9-like) IncN plasmids (publicly available; this study), and their geographic origin/dates of isolation. Plasmid sequence names in red are those from this study, derived from PacBio data unless otherwise specified. Aligned bars adjacent to plasmid names represent plasmid sequences: light grey denotes regions with 100% sequence identity; black represents nucleotide diversity between sequences; and thin lines represent indels. Coding sequences are represented by fat arrows below individual sequence bars and are colour-coded as per the colour key. The schematic depicts examples of evolutionary events described in the text: (a) single nucleotide level change, (b) small indels (≤100bp), (c) large indels (>100bp), (d) recombination events.

In Plasmid-12 (FJ223605), Tn*4401*b has inserted into a hybrid Tn*2*-Tn*3*-like element (with associated drug resistance genes including *bla*_TEM-1_, *bla*_OXA-9_, and several aminoglycoside resistance genes), albeit in the absence of target sequence duplication, possibly as the result of an intra-molecular, replicative transposition event generating mismatched target site sequences (L TSS = TATTA; R TSS = GTTCT). This complex is in turn located between two IS*15DIV* (IS15Δ)/IS*26*-like elements flanked by 8bp inverted repeats, and located between the *traI* (891bp from 3’ end) and *pld* loci (~28Kb; Figure 3A). The backbone components of the IncN Plasmid-12 are consistent with those seen in an NIH outbreak^8^ and in a rearranged version in a University of Virginia outbreak (CAV1043; 2008)^7^. From this study, plasmids from ecol_224, ecol_881, ecol_AZ159, ecol_422, and scaffolds from ecol_AZ151, ecol_744, ecol_AZ150 all share near identical structures to Plasmid-12, with clustered nucleotide level variation present in the *traJ-traI* genes, consistent with a homologous recombination event affecting this region, and evidence of sporadic insertion/deletion events (Figure 3A). However, the *bla*_KPC-_Tn*4401* structures in these isolates are almost entirely degraded by the presence of other mobile genetic elements (MGEs), including Tn*2*/Tn*3*-like elements, IS*Kpn8/27* and Tn*1721.* In ecol_224, *bla*_KPC-2_ has been inserted into the IncN backbone as part of two repeat, inverted Tn*3*-like structures, flanked by a TTGCT TSD, and closer to *traI* (136bp from 3’ end) than the aforementioned *IS15DIV* (IS15Δ)/IS26-like complex in Plasmid-12 (Figure 3B). Although it is not possible to accurately trace the evolutionary history of this genomic region given the available data, the presence of shared signatures of this structure in ecol_422, ecol_744, ecol_881, ecol_AZ159, ecol_AZ150 and ecol_AZ151 suggest a shared acquisition, and multiple subsequent rearrangements mediated by the presence of the large number of MGEs flanking *bla*_KPC-2._

**Figure 3.**
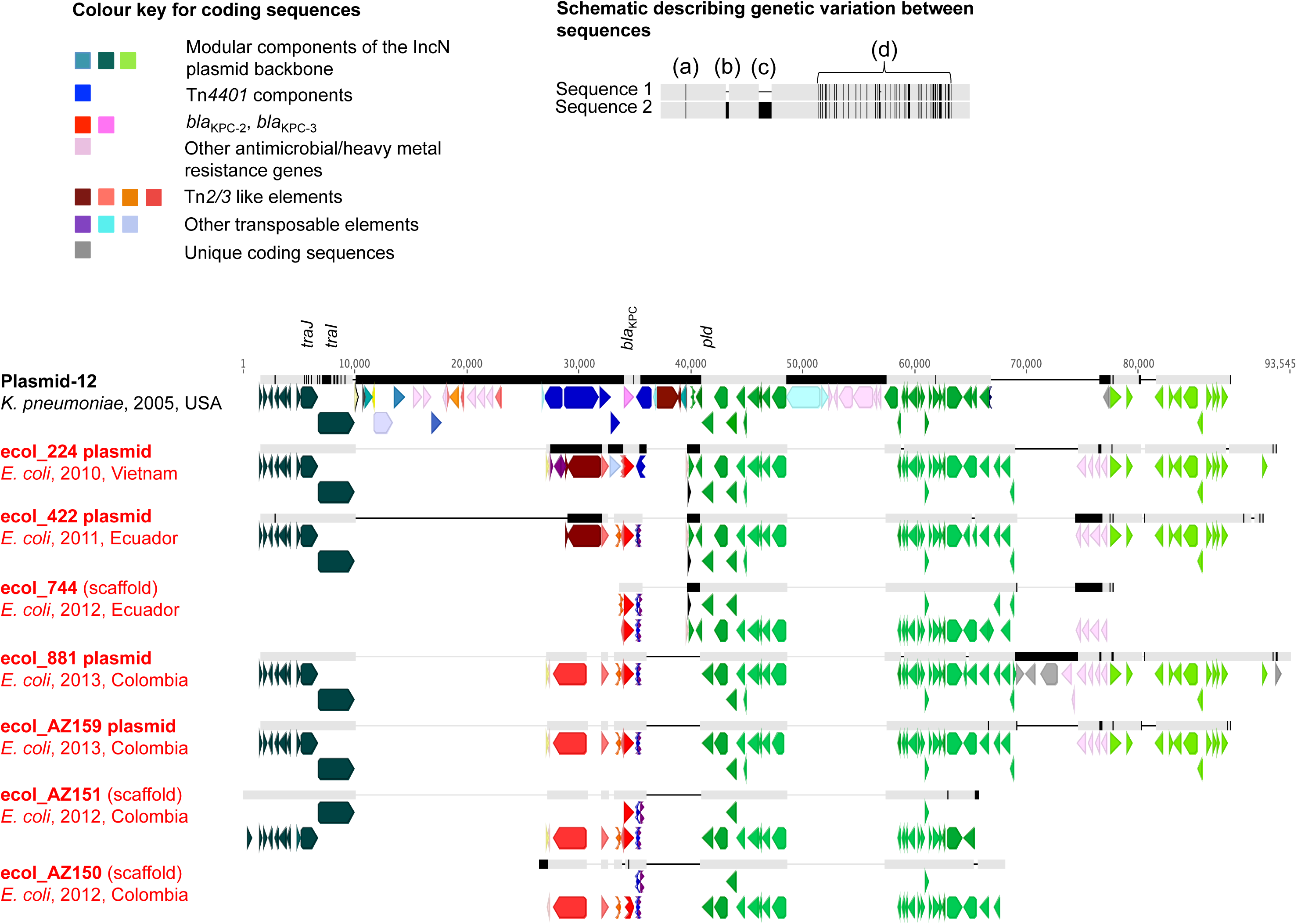
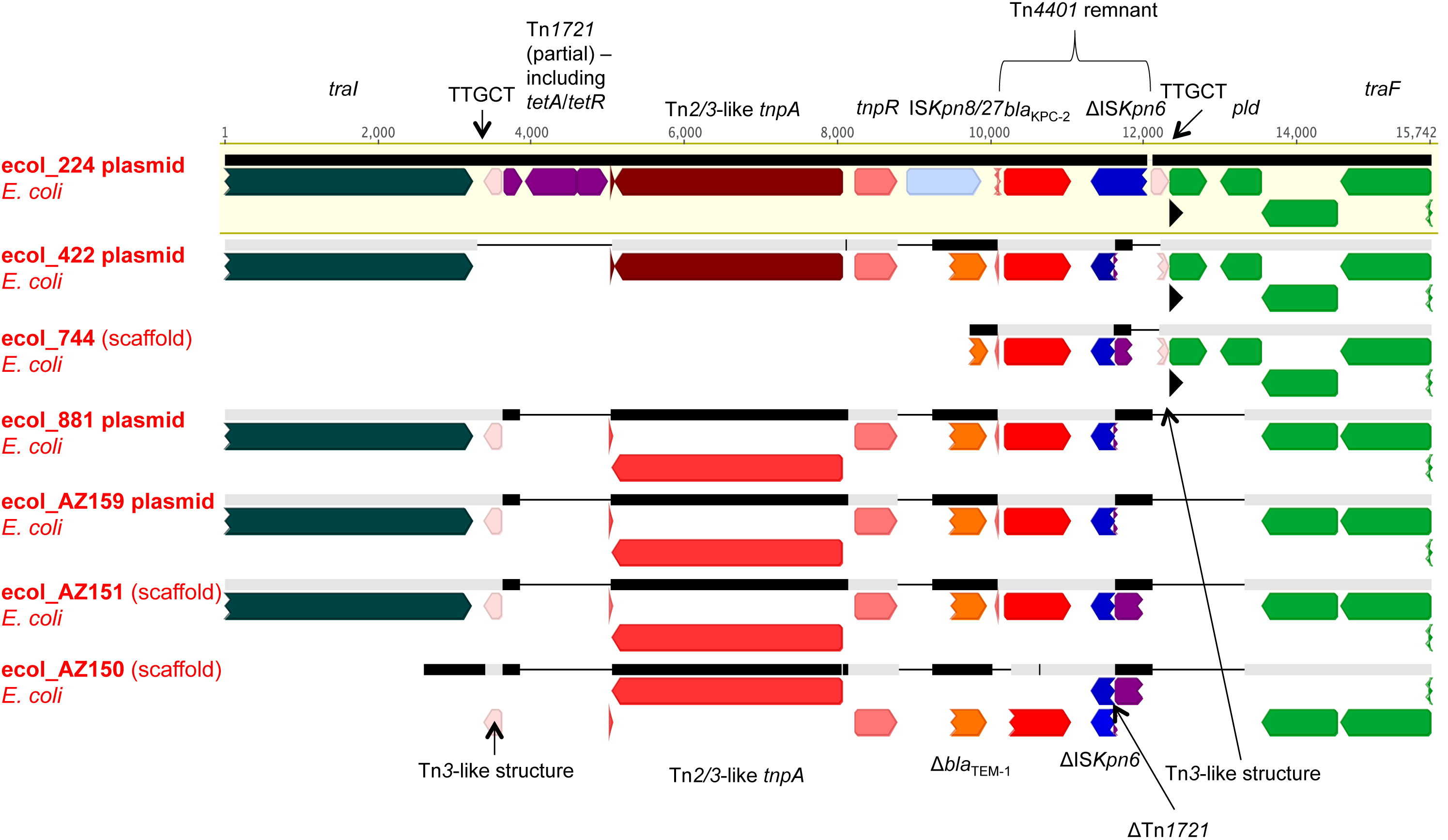
Comparison of FJ223605-like (Plasmid-12-like) IncN KPC plasmids from this study. Panel 3A. Geographic origin, dates of isolation and overall alignment of plasmid/scaffold structures. Plasmid sequence names in red are those from this study, derived from PacBio data unless denoted as “(scaffold)”, in which case these represent incomplete contigs from Illumina data. Aligned bars adjacent to plasmid names represent plasmid sequences: light grey denotes regions with 100% sequence homology;black represents nucleotide diversity between sequences; and thin lines represent indels. Coding sequences are represented by fat arrows below individual sequence bars and are colour-coded as per the colour key. The schematic depicts examples of evolutionary events described in the text:(a)single nucleotide level change, b) small indels (≤100bp), (c)large indels(>100bp), (d)recombination events. Panel 3B. Close-up of the region between *tral* and *pld* containing *bla*_KPC-2_ in study isolates. Coding sequences are colour-coded as in Figure 3A; sequence regions referred to in the text are annotated.

### Col-like plasmids may represent an important vector of transmission for *bla*_KPC_ in *E. coli*

Small col-like plasmids were the second most common type of plasmid carrying *bla*_KPC_ in *E. coli* (n=5 [plasmids with single replicons only]), but three of these were identical (*bla*_KPC-2_, 16,559bp), all isolated in Pittsburgh, USA, from ST131 isolates across a two year timeframe (ecol_276 [PacBio; 2010], ecol_356 [2011], ecol_875 [2013]). These three isolates additionally contained FIA, FIB, FII, X3 and X4 replicons, suggesting stable persistence of a clonal strain+plasmids over time, consistent with both SNV/core and accessory genome analyses (Figure 1, Figure S1).

The other two col-like plasmids effectively represent short stretches of DNA encoding different mobilization genes *(mbeA/mbeC/mbeD)* harnessed to Tn*4401/bla*_KPC_ modules. The 5bp sequences flanking Tn*4401* were consistent with direct, intermolecular transposition in both cases (ecol_870: TGTTT-TGTTT; ecol_867: TGTGA-TGTGA). A col/ *repA* co-integrate plasmid was also observed in this dataset (ecol_AZ161), in which Tn*4401*b was inserted between colE3 signature sequences and a Tn*3* element (Tn*4401* TSS: AGATA-GTTCT). The formation of such cointegrate plasmid structures in *E. coli* has also been previously described^34^, including that of a fused col/pKpQIL-like plasmid structure (pKpQIL being historically associated with *bla*_KPC_)^35^.

Col-like plasmids have been associated with KPC-producers in other smaller, regional studies^19,36^. Of concern, these small vectors have been shown to be responsible for the inter-species diffusion of *qnr* genes mediating fluoroquinolone resistance, even in the absence of any obvious antimicrobial selection pressure^37^. The significant association of col-like plasmids with particular *E. coli* STs (predominantly ST131) in this study could be one explanation for the disproportionate representation of *bla*_KPC_ in this lineage.

### Diverse Tn*4401* 5bp target site sequences (TSSs) support high transposon mobility

Complete Tn*4401* isoforms flanking *bla*_KPC-2_ or *bla*_KPC-3_ were observed in only 24/43 (56%) isolates, including Tn*4401*a/a-like (n=10; one isolate with a contig break upstream of *bla*_KPC_), Tn*4401*b (n=12), and Tn*4401*d (n=2) variants. Eleven different 5bp target site sequence (TSS) pairs were identified, of which 7 (64%) were not observed in any comparison plasmid downloaded from GenBank (Supplementary table 5). Tn*4401*a had three different 5bp TSSs, Tn*4401*b seven, and Tn*4401*d one. Most represented TSDs, but in three cases different 5bp TSSs were flanking Tn*4401*, consistent with both direct inter-and replicative intra-molecular transposition events.

From the full set of Genbank plasmids and *in vitro* transposition experiments carried out by others, 30 different types of 5bp TSS pairs have been characterized, seven in the experimental setting only^38^. The downloaded plasmids come from a range of species and time-points (2005-2014), although they may under-represent wider Tn*4401* insertion site diversity as a result of sampling biases. Our data however would be consistent with significant Tn*4401* mobility within *E. coli* following acquisition of diverse Tn*4401* isoforms and/or represent multiple importation events into *E. coli* from other species.

### The traditional association of *bla*_KPC_ with Tn4401 has been significantly eroded in KPC plasmids in *E. coli*

Notably, in the other 19/43 (44%) isolates the Tn*4401* structure had been degraded through replacement with MGEs, only some of which have been previously described^39,40^. Two isolates had novel Tn*4401*Δb structures (upstream truncations by IS*26* [ecol_270] or IS*26*-ΔIS*5075* [ecol_584]). A Tn*4401*e-like structure (255bp deletion upstream of *bla*_KPC_) was present in three isolates (ecol_227, ecol_316, ecol_583): this was further characterized in one complete PacBio plasmid assembly (ecol_316) and represented a rearrangement at the site of the L TSS of the ISK*pn7* element. In this plasmid, a second, partial Tn*4401* element was present without *bla*_KPC_, which would be consistent with an incomplete, replicative, intra-molecular transposition event (GGGAA = L TSS and R TSS on the two Tn*4401*b elements, in reverse orientation). Other motifs flanking *bla*_KPC_ included: hybrid Tn*2*/Tn*3* elements-ISK*pn8/27-bla*_KPC_ (n=1; ecol_224); IS*26*-Δ*tnpR(Tn3)-ISKpn8/27- bla*_KPC_-ΔTn*1721*-IS*26* (n=5; ecol_AZ153-AZ155, ecol_AZ166, ecol_AZ167); IS*Apu2-tnpR(Tn3)-* Δbla_TEM_ *-bla*_KPC_*- korC-klcA-*ΔTn*1721-*IS*26* (n=1; ecol_542); IS*26*-*tnpR*(Tn*3*)-Δbla_TEM_-*bla*_KPC_-korC-IS*26* (n=1; ecol_545); hybrid Tn*2*/Tn*3* elements + Δbla_TEM_-*bla*_KPC_-ΔΤn1721 (n=2; ecol_744, ecol_422), Tn*3* elements-Δ*bla*_TEM_-*bla*_KPC_- ΔTn*1721* (n=4; ecol_881, ecol_AZ151, ecol_AZ159, ecol_AZ150) and ΔTn*3*-Δ-ΔIS*3000* (Tn*3*-like) (n=1; ecol_AZ152). We were unable to assess the flanking context of *bla*_KPC_ in ecol_452 due to limitations of the assembly.

This apparent diversity in independently MGEs around the *bla*_KPC_ gene is a major concern, as it extends the means by which *bla*_KPC_ can be mobilized. Interestingly, as observed previously^41^, all the degraded Tn*4401* sequences in this dataset were associated with variable stretches of flanking Tn*2*/*3*-like sequences, suggesting that the insertion of Tn*4401* into a Tn*2*/Tn*3*-like context may have enabled the latter to act as a hotspot for the insertion of other MGEs^7^. A particular worry is the association with IS*26*, which has been linked to the dissemination of several other resistance genes in *E. coli,* including CTX-M ESBLs^22,42^; is able to increase the expression of closely co-located resistance genes^43^; participates in co-integrate formation and hence plasmid rearrangement^44^; and enhances the occurrence of other IS*26*-mediated transfer events into plasmids harbouring IS*26*^44^.

## DISCUSSION

This study of an unselected set of KPC-*E*. *coli,* obtained from two global resistance surveillance schemes, has demonstrated that the genetic structures associated with *bla*_KPC_ are highly diverse, at all genetic levels, including: (i) host bacterial strain; (ii) plasmid types; (iii) associated transposable MGEs, including transposons and insertion sequences; and (iv) *bla*_KPC_ alleles. This has previously been observed within institutional, poly-species outbreaks, particularly for non-*E*. *coli* Enterobacteriaceae^7,8^, as well as in a more recent study of nine KPC-*E*. *coli* from the US^45^. We have identified evidence of global and regional spread at the strain and plasmid levels, including signatures consistent with inter-species spread of plasmids in both these geographic contexts, over short timeframes. Although the geographic reach of sampling has been more substantial than any other similar study to date, some limitations in the sampling consistency of both the SMART and Astra Zeneca surveillance schemes has been observed^20^ (e.g. isolates from China were only submitted to these schemes in 2008, 2012 and 2013).

We utilized long-read sequencing methods on only a subset of isolates, given resource limitations, which allowed us to completely resolve chromosomal and plasmid structures in less than half the isolates. Nevertheless, despite this drawback, we have still highlighted the extraordinary diversity amongst these strains. This study, along with other recent analyses utilizing long-read sequencing to fully close important antimicrobial resistance plasmid structures^7,8^, also demonstrates the difficulty in making adequate evolutionary comparisons between these structures, given the absence of any effective phylogenetic methods to characterize the genetic histories for these structures where rearrangements are common, and events are not restricted to single nucleotide mutations.

This study has demonstrated the particular association of *bla_KPC_* in *E. coli* with IncN plasmid structures, which have been associated with the spread of other antimicrobial resistance elements^46^, as well as col-like plasmids, which are small, potentially highly mobile, and generally high copy number units. It has also highlighted that the traditional association of *bla*_KPC_ with Tn*4401* has been eroded in *E. coli,* with the complete Tn*4401* structure absent in 50% of strains investigated. This finding is in contrast to the majority of global descriptions in *K. pneumoniae* where *bla*_KPC_ has been stably associated with largely intact Tn*4401* isoforms for more than a decade. Instead, multiple other shorter mobile units, such as Tn*2*/Tn*3*-like elements and IS*26*, now appear to be commonly involved in the dispersal of *bla*_KPC_ in *E. coli.* These MGEs have been associated with the spread of multiple resistance mechanisms, such as *bla*_TEM_ and *bla*_CTX-M_, and will potentially similarly contribute to the dispersal of *bla*_KPC_ in *E. coli.* We did not undertake any functional assays investigating the experimental dynamics of *bla*_KPC_ transmission in *E. coli* to support this hypothesis, but this would be illuminating and important work for future studies.

The global emergence and spread of *bla*_KPC_ in *E. coli* has been driven by multiple mechanisms, including local and international spread of highly genetically related strains, exchange of plasmids with other Enterobacteriaceae and between *E. coli* lineages, transposition events within the species, and a breakdown of the traditional association of *bla*_KPC_ with Tn*4401.* The genetic flexibility observed is impressive, and concerning, particularly given that only a reasonably small number of KPC-*E*. *coli* over a short timeframe were characterized.

The diversity observed in this study has major implications for both surveillance and the clinical epidemiology of *E. coli.* Tracking the spread of resistance genes in the context of such multi-level genetic variability is complicated, even with a high-resolution typing method such as whole genome sequencing. The association of *E. coli,* both a common pathogen and commensal in a wide range of environmental/animal reservoirs, with MGEs (col-like plasmids, IS*26)* that have been shown to facilitate the dissemination of other successful resistance genes even in the absence of antimicrobial selection pressures, may represent a difficult, if not impossible, situation to control.

## METHODS

### Isolate collection and sampling frames

Isolates were obtained from two global antimicrobial resistance surveillance schemes (The Merck Study for Monitoring Antimicrobial Resistance Trends [SMART], 2008-2012; AstraZeneca global surveillance study of antimicrobial resistance, 2012-2013; 417 institutions operating in 95 countries), as previously described^20^. Of 55,874 isolates collected, 45 (0.08%) were positive for *bla*_KPC_ by PCR (n=7 from 2010, 10 from 2011, 13 from 2012, 15 from 2013 – Supplementary table 1). Isolates had been previously characterized using partial, sequenced-based typing methods, including multi-locus sequence typing (MLST; Achtman scheme), *fimH* typing, PCR for beta-lactamases, strain/plasmid PFGE (Supplementary table 1) ^20^.

### DNA extraction, sequencing and sequence data processing

All isolates were sequenced on the Illumina MiSeq; a subset of 18 were purposively selected for PacBio sequencing, to represent a range of years of isolation, geographic location, standard ST, plasmid size and resistance gene content (based on laboratory typing). DNA for sequencing was extracted from sub-cultures of bacterial stocks (frozen at -80°C) using the Qiagen Genomic tip 100/G extraction kit, as per the manufacturer’s instructions (Qiagen, Hilden, Germany; catalogue no: 10243).

DNA libraries for MiSeq sequencing were generated and normalized using 300 base, paired-end Nextera XT DNA library preparation kits (Illumina, San Diego, CA, USA). PacBio sequencing on the subset of strains was performed as previously described ^47^; in these cases, the same DNA extract was used for both Illumina and PacBio sequencing approaches.

Short-read Illumina data was processed as previously described^22^. Core variable sites (base called in all sequences, excluding “N” or “-” calls) derived from mapping to the SE15 reference were “padded” with invariant sites in a proportion consistent with the GC content and length of the reference genome (4.72Mb, 51% average GC content), to generate a modified alignment of input sequences to generate phylogenies. These were done using RaxML (Version 7.7.6) ^48^, with a generalized time reversible model, four gamma categories (relative rates of mutation across categories), bootstrapped 100 times. De novo assemblies of short-read Illumina data were generated using the A5-MiSeq pipeline (version 20140604)^49^.

Plasmid and chromosome structures were closed by resolving repeats at the ends of assembled, polished, PacBio contigs. Illumina reads were mapped to the resulting assemblies using bwa-MEM version 0.7.9a-r786 with default settings^50^. Read pileups were visualized in Geneious^51^; mismatches between the sequence derived from mapping and the reference PacBio assemblies were inspected manually to identify the correct structure, resulting in a final consensus sequence used for subsequent analyses and submission to GenBank. Unmapped reads were de novo assembled using the A5-MiSeq pipeline 20140604^49^ to capture small plasmids that may have been filtered out due to size-selection of DNA fragments >7,000 bases implemented prior to PacBio sequencing.

All plasmid structures and de novo assemblies were annotated using PROKKA^52^, with subsequent manual refinement of annotations for regions of interest using BLASTn ^53^ and the NCBI bacterial and ISFinder databases ^54^. Alignments of sequence structures were visualized and modified in Geneious.

### Core/accessory genome comparisons

These were undertaken using the pangenome pipeline, ROARY ^25^, by inputting the *.gff files generated from the PROKKA annotation of each of the Illumina de novo assemblies (default settings). Comparisons were made separately for all isolates and the ST131 subset. The output gene_presence_absence.csv files were processed using the pheatmap function in R. Resistance genes were identified using ResisType, an inhouse tool [scripts available at: https://github.com/hangphan/resisType]. These were plotted on the maximum likelihood phylogenies using the Ape package in R.

### Comparisons with publicly available KPC plasmid sequences

All complete KPC RefSeq sequences available in GenBank in May 2015 were identified using the search terms “plasmid” + “KPC” + “complete sequence”. The resulting list was filtered manually to exclude any additional sequences present that were not complete plasmid sequences. In total 73 plasmid sequences were included (Supplementary Table 5).

For the IncN plasmid comparisons, we included the following from our dataset: (i) six cases where PacBio sequencing had fully resolved the *bla*_KPC_ IncN plasmid; (ii) two cases where Illumina sequencing had fully resolved the *bla*_KPC_ IncN plasmid; (iii) two cases where the IncN rep and *bla*_KPC_ were co-located on the same, incomplete contig; and (iv) two cases where *bla*_KPC_ present in isolates containing an IncN rep and on contigs that had similar plasmid backbones to the other IncN plasmids under scrutiny.

### Availability of Data and Materials

The data sets (Illumina raw reads, PacBio assemblies) supporting the results of this article are available in NCBI’s GenBank/SRA, under the project accession: PRJNA316786 (https://www.ncbi.nlm.nih.gov/bioproject/?term=316786).

## ACKNOWLEDGEMENTS

We acknowledge the contributions of the laboratory, healthcare and administrative teams contributing to the SMART and Astra Zeneca global antimicrobial resistance surveillance programs, and the Modernising Medical Microbiology Informatics Group (MMMIG). For this study, the MMMIG consisted of Adam Giess, Carlos Del Ojo Elias, Milind Acharya, Nicholas Sanderson, Trien Do and Vasiliki Kostiou.

## AUTHOR CONTRIBUTIONS

NS and JP conceived of the study. Significant contributions to sample collection, laboratory processing and sequencing were made by GP, LWA, LP, PB, MRM, NS and JP. Short-read (Illumina) sequencing was performed by LWA and LP; long-read (PacBio) sequencing by RS and AK. Sequence data processing and analysis were performed by AES, HTTP and NS. NS drafted the manuscript, which was reviewed and improved by all authors, including ASW, TEAP, DWC and AJM.

## ADDITIONAL INFORMATION

### FUNDING INFORMATION

NS is currently funded through a Public Health England/University of Oxford Clinical Lectureship; the sequencing work was also partly funded through a previous Wellcome Trust Doctoral Research Fellowship (#099423/Z/12/Z). Additional funding support was provided by a research grant from Calgary Laboratory Services (#10006465), and by the Health Innovation Challenge Fund (a parallel funding partnership between the Wellcome Trust [WT098615/Z/12/Z] and the Department of Health [grant HICF-T5-358]). This research was supported by the National Institute for Health Research (NIHR) Oxford Biomedical Research Center (BRC) Program, and the Health Protection Research Unit (NIHR HPRU) in Healthcare Associated Infections and Antimicrobial Resistance at the University of Oxford, in partnership with Public Health England (PHE).

The funders had no role in study design, data collection and interpretation, or the decision to submit the work for publication. The views expressed are those of the author(s) and not necessarily those of the NHS, the NIHR or the Department of Health.

## Competing interest

The authors declare that they have no competing interests.

## REFERENCES

1 Nordmann, P., Naas, T. & Poirel, L. Global spread of Carbapenemase-producing Enterobacteriaceae. Emerg Infect Dis 17, 1791-1798, doi:10.3201/eid1710.110655 (2011).

2 Doi, Y. & Paterson, D. L. Carbapenemase-producing Enterobacteriaceae. Semin Respir Crit Care Med 36, 74-84, doi:10.1055/s-0035-1544208 (2015).

3 Nordmann, P. & Poirel, L. The difficult-to-control spread of carbapenemase producers among Enterobacteriaceae worldwide. Clin Microbiol Infect 20, 821-830, doi:10.1111/1469-0691.12719 (2014).

4 Munoz-Price, L. S. et al. Clinical epidemiology of the global expansion of Klebsiella pneumoniae carbapenemases. Lancet Infect Dis 13, 785-796, doi:10.1016/S1473-3099(13)70190-7 (2013).

5 Nordmann, P., Cuzon, G. & Naas, T. The real threat of Klebsiella pneumoniae carbapenemase-producing bacteria. Lancet Infect Dis 9, 228-236, doi:10.1016/S1473-3099(09)70054-4 (2009).

6 Pitout, J. D., Nordmann, P. & Poirel, L. Carbapenemase-Producing Klebsiella pneumoniae, a Key Pathogen Set for Global Nosocomial Dominance. Antimicrob Agents Chemother 59, 5873-5884, doi:10.1128/AAC.01019-15 (2015).

7 Sheppard, A. E. et al. Nested Russian Doll-Like Genetic Mobility Drives Rapid Dissemination of the Carbapenem Resistance Gene blaKPC. Antimicrob Agents Chemother 60 3767-3778, doi:10.1128/AAC.00464-16 (2016).

8 Conlan, S. et al. Single-molecule sequencing to track plasmid diversity of hospital-associated carbapenemase-producing Enterobacteriaceae. Sci Transl Med 6, 254ra126, doi:10.1126/scitranslmed.3009845 (2014).

9 Deshpande, L. M., Rhomberg, P. R., Sader, H. S. & Jones, R. N. Emergence of serine carbapenemases (KPC and SME) among clinical strains of Enterobacteriaceae isolated in the United States Medical Centers: report from the MYSTIC Program (1999-2005). Diagn Microbiol Infect Dis 56, 367-372, doi:10.1016/j.diagmicrobio.2006.07.004 (2006).

10 Hong, T. et al. Escherichia coli: development of carbapenem resistance during therapy. Clin Infect Dis 40, e84-86, doi:10.1086/429822 (2005).

11 NavonVenezia, S. et al. Plasmid-mediated imipenem-hydrolyzing enzyme KPC-2 among multiple carbapenem-resistant Escherichia coli clones in Israel. Antimicrob Agents Chemother 50, 3098-3101, doi:10.1128/AAC.00438-06 (2006).

12 Mathers, A. J., Peirano, G. & Pitout, J. D. The role of epidemic resistance plasmids and international high-risk clones in the spread of multidrug-resistant Enterobacteriaceae. Clin Microbiol Rev 28, 565-591, doi:10.1128/CMR.00116-14 (2015).

13 Gona, F. et al. In vivo multiclonal transfer of bla(KPC-3) from Klebsiella pneumoniae to Escherichia coli in surgery patients Clin Microbiol Infect 20, O633-635, doi:10.1111/1469-0691.12577 (2014).

14 Luo, Y. et al. Characterization of KPC-2-producing Escherichia coli, Citrobacter freundii, Enterobacter cloacae, Enterobacter aerogenes, and Klebsiella oxytoca isolates from a Chinese Hospital. Microb Drug Resist 20, 264-269, doi:10.1089/mdr.2013.0150 (2014).

15 Robledo, I. E., Aquino, E. E. & Vazquez, G. J. Detection of the KPC gene in Escherichia coli, Klebsiella pneumoniae, Pseudomonas aeruginosa, and Acinetobacter baumannii during a PCR-based nosocomial surveillance study in Puerto Rico. Antimicrob Agents Chemother 55, 2968-2970, doi:10.1128/AAC.01633-10 (2011).

16 Mavroidi, A. et al. Emergence of Escherichia coli sequence type 410 (ST410) with KPC-2 beta-lactamase. Int J Antimicrob Agents 39, 247-250, doi:10.1016/j.ijantimicag.2011.11.003 (2012).

17 Xu, G., Jiang, Y., An, W., Wang, H. & Zhang, X. Emergence of KPC-2-producing Escherichia coli isolates in an urban river in Harbin, China. World J Microbiol Biotechnol 31, 1443-1450, doi:10.1007/s11274-015-1897-z (2015).

18 Cai, J. C., Zhang, R., Hu, Y. Y., Zhou, H. W. & Chen, G. X. Emergence of Escherichia coli sequence type 131 isolates producing KPC-2 carbapenemase in China. Antimicrob Agents Chemother 58, 1146-1152, doi:10.1128/AAC.00912-13 (2014).

19 O'Hara, J. A.et al. Molecular epidemiology of KPC-producing Escherichia coli: occurrence of ST131-fimH30 subclone harboring pKpQIL-like IncFIIk plasmid. Antimicrob Agents Chemother 58, 4234-4237, doi:10.1128/AAC.02182-13 (2014).

20 Peirano, G. et al. Global incidence of carbapenemase-producing Escherichia coli ST131. Emerg Infect Dis 20, 1928-1931, doi:10.3201/eid2011.141388 (2014).

21 Petty, N. K. et al. Global dissemination of a multidrug resistant Escherichia coli clone. Proc Natl Acad Sci U S A 111, 5694-5699, doi:10.1073/pnas.1322678111 (2014).

22 Stoesser, N. et al. Evolutionary History of the Global Emergence of the Escherichia coli Epidemic Clone ST131. MBio 7, doi:10.1128/mBio.02162-15 (2016).

23 Banerjee, R. & Johnson, J. R. A new clone sweeps clean: the enigmatic emergence of Escherichia coli sequence type 131. Antimicrob Agents Chemother 58, 4997-5004, doi:10.1128/AAC.02824-14 (2014).

24 Zhong, Y. M. et al. Emergence and spread of O16-ST131 and O25b-ST131 clones among faecal CTX-M-producing Escherichia coli in healthy individuals in Hunan Province, China. J Antimicrob Chemother 70, 2223-2227, doi:10.1093/jac/dkv114 (2015).

25 Page, A. J. et al.Roary: rapid large-scale prokaryote pan genome analysis. Bioinformatics 31, 3691-3693, doi:10.1093/bioinformatics/btv421 (2015).

26 Touchon, M. et al. Organised genome dynamics in the Escherichia coli species results in highly diverse adaptive paths. PLoS Genet 5, e1000344, doi:10.1371/journal.pgen.1000344 (2009).

27 Martinez, T., Vazquez, G. J., Aquino, E. E., Martinez, I. & Robledo, I. E. ISEcp1-mediated transposition of blaKPC into the chromosome of a clinical isolate of Acinetobacter baumannii from Puerto Rico. J Med Microbiol 63, 1644-1648, doi:10.1099/jmm.0.080721-0 (2014).

28 Chen, L.et al. Genome Sequence of a Klebsiella pneumoniae Sequence Type 258 Isolate with Prophage-Encoded K. pneumoniae Carbapenemase. Genome Announc 3, doi:10.1128/genomeA.00659-15 (2015).

29 Norman, A., Hansen, L. H. & Sorensen, S. J. Conjugative plasmids: vessels of the communal gene pool. Philos Trans R Soc Lond B Biol Sci 364, 2275-2289, doi:10.1098/rstb.2009.0037 (2009).

30 Watve, M. M., Dahanukar, N. & Watve, M. G. Sociobiological control of plasmid copy number in bacteria. PLoS One 5, e9328, doi:10.1371/journal.pone.0009328 (2010).

31 Paulsson, J. Multileveled selection on plasmid replication. Genetics 161, 1373-1384 (2002).

32 San Millan, A. et al. Small-plasmid-mediated antibiotic resistance is enhanced by increases in plasmid copy number and bacterial fitness. Antimicrob Agents Chemother 59, 3335-3341, doi:10.1128/AAC.00235-15 (2015).

33 Gootz, T. D.et al. Genetic organization of transposase regions surrounding blaKPC carbapenemase genes on plasmids from Klebsiella strains isolated in a New York City hospital. Antimicrob Agents Chemother 53 1998-2004, doi:10.1128/AAC.01355-08 (2009).

34 Peterson, B. C., Hashimoto, H. & Rownd, R. H. Cointegrate formation between homologous plasmids in Escherichia coli. J Bacteriol 151, 1086-1094 (1982).

35 Villa, L.et al.Reversion to susceptibility of a carbapenem-resistant clinical isolate of Klebsiella pneumoniae producing KPC-3. J Antimicrob Chemother 68, 2482-2486, doi:10.1093/jac/dkt235 (2013)).

36 Partridge, S. R. et al.Emergence of blaKPC carbapenemase genes in Australia. Int J Antimicrob Agents 45, 130-136, doi:10.1016/j.ijantimicag.2014.10.006 (2015).

37 Pallecchi, L.et al.Small qnrB-harbouring ColE-like plasmids widespread in commensal enterobacteria from a remote Amazonas population not exposed to antibiotics. J Antimicrob Chemother 66, 1176-1178, doi:10.1093/jac/dkr026 (2011).

38 Cuzon, G., Naas, T. & Nordmann, P. Functional characterization of Tn4401, a Tn3-based transposon involved in blaKPC gene mobilization. Antimicrob Agents Chemother 55, 5370-5373, doi:10.1128/AAC.05202-11 (2011).

39 Chen, L. et al. Carbapenemase-producing Klebsiella pneumoniae: molecular and genetic decoding. Trends Microbiol 22, 686-696, doi:10.1016/j.tim.2014.09.003 (2014).

40 Shen, P. et al.Novel genetic environment of the carbapenem-hydrolyzing beta-lactamase KPC-2 among Enterobacteriaceae in China. Antimicrob Agents Chemother 53, 4333-4338, doi:10.1128/AAC.00260-09 (2009).

41 Liu, L. et al. blaCTX-M-1/9/1 Hybrid Genes May Have Been Generated from blaCTX-M-15 on an IncI2 Plasmid. Antimicrob Agents Chemother 59, 4464-4470, doi:10.1128/AAC.00501-15 (2015).

42 Partridge, S. R., Zong, Z. & Iredell, J. R. Recombination in IS26 and Tn2 in the evolution of multiresistance regions carrying blaCTX-M-15 on conjugative IncF plasmids from Escherichia coli. Antimicrob Agents Chemother 55, 4971-4978, doi:10.1128/AAC.00025-11 (2011).

43 Lee, K. Y., Hopkins, J. D. & Syvanen, M. Direct involvement of IS26 in an antibiotic resistance operon. J Bacteriol 172, 3229-3236 (1990).

44 Harmer, C. J., Moran, R. A. & Hall, R. M. Movement of IS26-associated antibiotic resistance genes occurs via a translocatable unit that includes a single IS26 and preferentially inserts adjacent to another IS26. MBio 5, e01801-01814, doi:10.1128/mBio.01801-14 (2014).

45 Chavda, K. D., Chen, L., Jacobs, M. R., Bonomo, R. A. & Kreiswirth, B. N. Molecular Diversity and Plasmid Analysis of KPC-Producing Escherichia coli. Antimicrob Agents Chemother 60, 4073-4081, doi:10.1128/AAC.00452-16 (2016).

46 Garcia-Fernandez, A.et al.Multilocus sequence typing of IncN plasmids. J Antimicrob Chemother 66, 1987-1991, doi:10.1093/jac/dkr225 (2011).

47 Mathers, A. J. et al. Klebsiella pneumoniae carbapenemase (KPC)-producing K. pneumoniae at a single institution: insights into endemicity from whole-genome sequencing. Antimicrob Agents Chemother 59, 1656-1663, doi:10.1128/AAC.04292-14 (2015).

48 Stamatakis, A. RAxML version 8: a tool for phylogenetic analysis and postanalysis of large phylogenies. Bioinformatics 30, 1312-1313, doi:10.1093/bioinformatics/btu033 (2014).

49 Coil, D., Jospin, G. & Darling, A. E. A5-miseq: an updated pipeline to assemble microbial genomes from Illumina MiSeq data. Bioinformatics 31, 587-589, doi:10.1093/bioinformatics/btu661 (2015).

50 Li, H. & Durbin, R. Fast and accurate short read alignment with BurrowsWheeler transform. Bioinformatics 25, 1754-1760, doi:10.1093/bioinformatics/btp324 (2009).

51 Kearse, M. M. R., Wilson, A.; Stones-Havas, S.; Cheung, M.; Sturrock, S.; Buxton, A.; Markowitz, S.; Duran, C.; Thierer, T.; Ashton, B.; Metjies, P.; Drummond A. Geneious Basic: an integrated and extendable desktop software platform for the organization and analysis of sequence data. Bioinformatics 28, 1647-1649 (2012).

52 Seemann, T. Prokka: rapid prokaryotic genome annotation. Bioinformatics 30, 2068-2069, doi:10.1093/bioinformatics/btu153 (2014).

53 Altschul, S. F., Gish, W., Miller, W., Myers, E. W. & Lipman, D. J. Basic local alignment search tool. J Mol Biol 215, 403-410, doi:10.1016/S0022-2836(05)80360-2 (1990).

54 Siguier, P., Perochon, J., Lestrade, L., Mahillon, J. & Chandler, M. ISfinder: the reference centre for bacterial insertion sequences. Nucleic Acids Res 34, D32-36, doi:10.1093/nar/gkj014 (2006).

55 Carattoli, A.et al.In silico detection and typing of plasmids using PlasmidFinder and plasmid multilocus sequence typing. Antimicrob Agents Chemother 58, 3895-3903, doi:10.1128/AAC.02412-14 (2014).

